# Cell-intrinsic activation of Orai1 regulates human T cell motility

**DOI:** 10.1101/128595

**Authors:** Tobias X. Dong, Milton L. Greenberg, Sabrina Leverrier, Ying Yu, Ian Parker, Joseph L. Dynes, Michael D. Cahalan

## Abstract

Ca^2+^ signaling through the store-operated Ca^2+^ channel, Orai1, is crucial for T cell function, but a role in regulating T cell motility in lymph nodes has not been previously reported. Tracking human T cells in immunodeficient mouse lymph nodes and in microfabricated PDMS channels, we show that inhibition of Orai1 channel activity with a dominant-negative Orai1-E106A construct increases average T cell velocities by reducing the frequency of pauses in motile T cells. Orai1-dependent motility arrest occurs spontaneously during confined motility *in vitro*, even in the absence of extrinsic cell contacts or antigen recognition. Utilizing a novel ratiometric genetically encoded cytosolic Ca^2+^ indicator, Salsa6f, we show these spontaneous pauses during T cell motility *in vitro* coincide with episodes of spontaneous cytosolic Ca^2+^ signaling. Our results demonstrate that Orai1, activated in a cell-intrinsic manner, regulates T cell motility patterns that accompany immune surveillance.

---

To initiate the adaptive immune response, T cells must make direct contact with antigen-presenting cells (APCs) in the lymph node, enabling T cell receptors (TCRs) to engage peptide-bound MHC molecules presented on the APC surface. Because cognate antigens are rare for any given TCR, many APCs must be scanned to identify those bearing cognate antigens. Thus, optimizing T cell motility to balance search sensitivity and speed is crucial for efficient antigen search and proper immune function (Cahalan and Parker 2005, Krummel, Bartumeus et al. 2016). Both cell-intrinsic and environmental factors have been proposed to regulate T cell motility within lymph nodes and peripheral tissues (Miller, Wei et al. 2002, Bousso and Robey 2003, Mempel, Henrickson et al. 2004, Mrass, Petravic et al. 2010). T cell motility in lymph nodes, sometimes referred to as “basal motility”, has been likened to diffusive Brownian motion, resembling a “stop-and-go” random walk with an overall exploratory spread that results in a linear mean squared displacement over time (Miller, Wei et al. 2002). Subsequent studies defined a role of cellular cues in guiding T cell migration, such as contact with the lymph node stromal cell network or short-term encounters with resident dendritic cells (Miller, Hejazi et al. 2004, Bajenoff, Egen et al. 2006, Khan, Headley et al. 2011). Whereas the basic signaling mechanisms for cell-intrinsic induction of random motility have been previously explored in several cell types (Petrie, Doyle et al. 2009), it remains unclear if such mechanisms apply in T cells.

Upon T cell recognition of cognate antigen, TCR engagement results in an elevated cytosolic Ca^2+^ concentration that acts as a “STOP” signal to halt motility and anchor the T cell to the site of antigen presentation (Donnadieu, Bismuth et al. 1994, Negulescu, Krasieva et al. 1996, Dustin, Bromley et al. 1997, Bhakta, Oh et al. 2005, Moreau, Lemaitre et al. 2015). The predominant mechanism for increasing cytosolic Ca^2+^ in T cells is through store-operated Ca^2+^ entry (SOCE), which is mediated by the molecular components STIM1 and Orai1. TCR stimulation triggers depletion of intracellular Ca^2+^ stores in the endoplasmic reticulum (ER), resulting in translocation of the ER-resident Ca^2+^ sensor STIM1 to specialized ER-plasma membrane (PM) junctions where Orai1 channels aggregate into puncta and activate to allow sustained Ca^2+^ influx (Liou, Kim et al. 2005, Roos, DiGregorio et al. 2005, Zhang, Yu et al. 2005, Luik, Wu et al. 2006, Vig, Beck et al. 2006, Zhang SL 2006, Calloway, Vig et al. 2009, Wu, Covington et al. 2014). Orai1 channel activity is crucial for immune function, as human mutations in Orai1 result in severe combined immunodeficiency (SCID) (Feske, Gwack et al. 2006). Additional roles of Orai1 have been defined in chemotaxis to certain chemokines and T cell homing to lymph nodes (Greenberg, Yu et al. 2013); actin cytoskeleton rearrangement (Schaff, Dixit et al. 2010, Dixit, Yamayoshi et al. 2011, Babich and Burkhardt 2013); migration during shear flow (Schaff, Dixit et al. 2010, Dixit, Yamayoshi et al. 2011); lipid metabolism (Maus, Cuk et al. 2017); and dendritic spine maturation in neurons (Korkotian, Oni-Biton et al. 2017). However, despite their contributions to other aspects of T cell function, no role has been identified for Orai1 channels in T cell motility patterns underlying scanning behavior.

Ca^2+^ imaging studies commonly rely on small synthetic Ca^2+^ indicators such as the ratiometric probe fura-2. However, synthetic indicators must be loaded into cells and cannot be targeted to subcellular compartments or specific cell populations. Furthermore, synthetic indicators slowly leak out of cells over time, and thus are unsuitable for long-term studies. To overcome these limitations, genetically encoded Ca^2+^ indicators (GECIs) were first developed two decades ago as probes based on Förster Resonance Energy Transfer (FRET) (Miyawaki, Llopis et al. 1997, Romoser, Hinkle et al. 1997, Perez Koldenkova and Nagai 2013). Since then, GECIs have undergone remarkable improvements in variety and functionality, including the optimization of single fluorescent protein-based indicators such as the GCaMP and the GECO series (Nakai, Ohkura et al. 2001, Zhao, Araki et al. 2011), as well as the new FRET-based TN indicators utilizing troponin C instead of calmodulin for the Ca^2+^ sensing element (Heim and Griesbeck 2004, Thestrup, Litzlbauer et al. 2014). Whereas single fluorescent protein-based indicators have high brightness and fast response kinetics, as non-ratiometric probes they are problematic for Ca^2+^ imaging in motile cells because fluorescence changes resulting from movement are indistinguishable from actual changes in Ca^2+^ levels. Here, we introduce a novel genetically encoded Ca^2+^ indicator, Salsa6f, consisting of GCaMP6f fused to tdTomato, providing high dynamic range and ratiometric capabilities.

In this study, we use human T cells to assess the role of Orai1 in cell motility. Expression of a dominant-negative Orai1-E106A construct was used to block Orai1 channel activity both *in vivo* within immunodeficient mouse lymph nodes (Greenberg, Yu et al. 2013), and *in vitro* within microfabricated polydimethylsiloxane (PDMS) chambers (Jacobelli, Friedman et al. 2010). We then utilize our genetically encoded Salsa6f probe to monitor spontaneous Ca^2+^ signaling in confined microchannels *in vitro*. Our results indicate that Ca^2+^ influx through Orai1 channels, activated intermittently in a cell-intrinsic manner, triggers spontaneous pauses during T cell motility that fine-tunes the search for cognate antigens.

## Results

### Validating a dominant-negative construct to inhibit Orai1 in human T cells

To study the role of Orai1 channel activity in T cell motility, we took a molecular genetic approach using the dominant-negative mutant Orai1-E106A to selectively eliminate ion conduction through the Orai1 pore. The glutamate residue at position 106 in human Orai1 forms the selectivity filter of the Orai1 pore (Prakriya, Feske et al. 2006, Vig, Beck et al. 2006, Yeromin, Zhang et al. 2006), and because the Orai1 channel is a functional hexamer (Hou, Pedi et al. 2012), mutation of E106 to neutrally charged alanine completely inhibits Ca^2+^ permeation in a potent dominant-negative manner (Greenberg, Yu et al. 2013). We confirmed Orai1 channel block by E106A by fura-2 Ca^2+^ imaging in activated human T cells transfected with either eGFP-tagged Orai1-E106A or empty vector for control. Following thapsigargin treatment to deplete the ER Ca^2+^ store, Ca^2+^ entry was greatly diminished in cells expressing eGFP-Orai1-E106A identified by fluorescence, referred to here as eGFP-E106^hi^ T cells, compared to empty vector-transfected control cells (Figure 1A). Ca^2+^ entry was also partially inhibited in some transfected T cells even though their eGFP fluorescence was too low to detect. We refer to this mixed population as eGFP-E106A^lo^ cells. To further confirm that eGFP-E106A inhibits T cell activation, we challenged transfected human T cells with autologous dendritic cells pulsed with the superantigen Staphylococcal enterotoxin B (Lioudyno, Kozak et al. 2008). T cell proliferation was markedly suppressed in eGFP-E106A^hi^ CD4^+^ and CD8^+^ T cells, but not in eGFP-E106A^lo^ T cells (Supplementary Figure 1A). This shows that reduced Orai1 channel activity present in eGFP-E106A^lo^ T cells is sufficient for T cell activation and proliferation. Taken together, these experiments show that eGFP-tagged Orai1-E106A expression can serve as a robust tool to assess cellular roles of Orai1 channel activity, and that transfected cells without detectable eGFP fluorescence can be used as an internal control.

**Figure 1.**
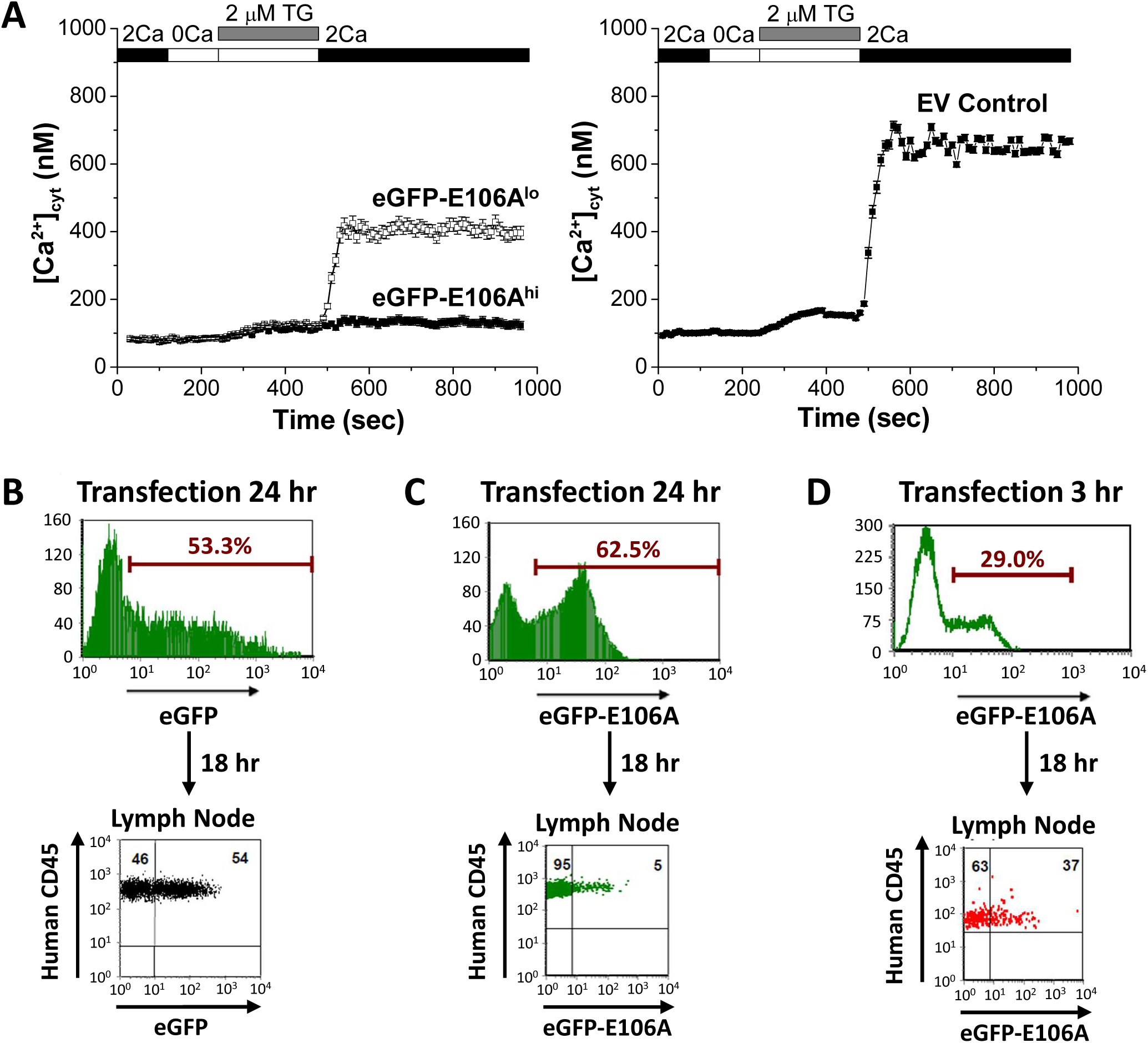
Effects of expressing Orai1-E106A on human T cell function. (**A**) Averaged thapsigargin-induced Ca^2+^ entry, measured by fura-2, in activated human CD4^+^ T cells transfected with eGFP-Orai1-E106A (left) or empty vector control (EV, right, n = 133 cells); eGFP-E106A transfected cells were grouped into two populations, either eGFP-E106A^hi^ with high eGFP fluorescence (solid squares, n = 43 cells) or eGFP-E106A^lo^ with no detectable eGFP fluorescence (empty squares, n = 115 cells); bars represent SEM, data representative of at least three different experiments. (**B**) Human CD3^+^ T cells transfected with eGFP for control and expression level was measured at 24 hr post-transfection before adoptive transfer into reconstituted NOD.SCI.β2 mice; cells were recovered from lymph nodes 18 hr later and eGFP fluorescence was used to measure homing to lymph nodes. (**C,D**) Human CD3^+^ T cells were transfected with eGFP-E106A and expression level was measured before adoptive transfer into reconstituted NOD.SCI.β2 mice either 24 hr (**C**) or 3 hr (**D**) post-transfection; cells were recovered from lymph nodes 18 hr later and eGFP fluorescence was used to measure homing to lymph nodes; data representative of independent experiments from 12 different donors.

Orai1 function in human T cell motility was evaluated *in vivo* using a human xenograft model in which immunodeficient NOD.SCID. β2 mice were reconstituted with human peripheral blood lymphocytes, followed by imaging of excised lymph nodes using two-photon microscopy (Greenberg, Yu et al. 2013). Reconstitution has been shown to produce a high density of human immune cells within the lymph nodes of immunodeficient mice (Mosier, Gulizia et al. 1988), simulating the crowded migratory environment experienced by T cells under normal physiological conditions. Three weeks after reconstitution, human T cells were purified from the same donor, transfected, and adoptively transferred into the reconstituted NOD.SCID.β2 mice (Supplementary Figure 1B). Whereas control T cells transfected with eGFP showed robust expression and successfully homed to lymph nodes following adoptive transfer 24 hr post-transfection (Figure 1B), eGFP-E106A transfected T cells did not home to lymph nodes when adoptively transferred 24 hr post-transfection (Figure 1C). This result confirms our previous study indicating that functional Orai1 channel activity is required for T cell homing to lymph nodes (Greenberg, Yu et al. 2013). To circumvent the homing defect, we injected eGFP-E106A transfected T cells only 3 hr post-transfection, before the expression level of eGFP-E106A had become sufficiently high to block lymph node entry (Figure 1D).

### Orai1 block increases human T cell motility within intact lymph node

To evaluate Orai1 function in T cell motility, we imaged human T cells within intact lymph nodes of reconstituted NOD.SCID.β2 mice by two-photon microscopy (Figure 2A). We found that eGFP-E106A^hi^ T cells migrated with higher average velocities than co-transferred, mock-transfected CMTMR-labeled T cells (12.8 ± 0.5 μm/min vs. 11.1 ± 0.5 μm/min, p = 0.0268; Figure 2B). Although both populations had similar maximum (Hodges-Lehmann median difference of -0.21 μm/min, -2.82 to 2.16 μm/min 95%CI) and minimum (Hodges-Lehmann median difference of -0.36 μm/min, -1.02 to 0.36 μm/min 95%CI) instantaneous cell velocities (Figure 2C), the arrest coefficient, defined by the fraction of time that cell velocity was < 2 μm/min, was six-fold lower for eGFP-E106A^hi^ T cells than for control T cells (0.02 ± 0.01 vs. 0.12 ± 0.03, p = 0.0406; Figure 2D). These differences in motility suggest that the increase in average cell velocity caused by Orai1 block is not due to eGFP-E106A^hi^ T cells moving faster than control T cells, but rather due to a reduced frequency of pausing. Consistent with this interpretation, no eGFP-E106A^hi^ T cells with average velocities < 7 μm/min were observed, unlike control T cells in which 23% of average velocities were < 7 µm/min (Figure 2E).

**Figure 2.**
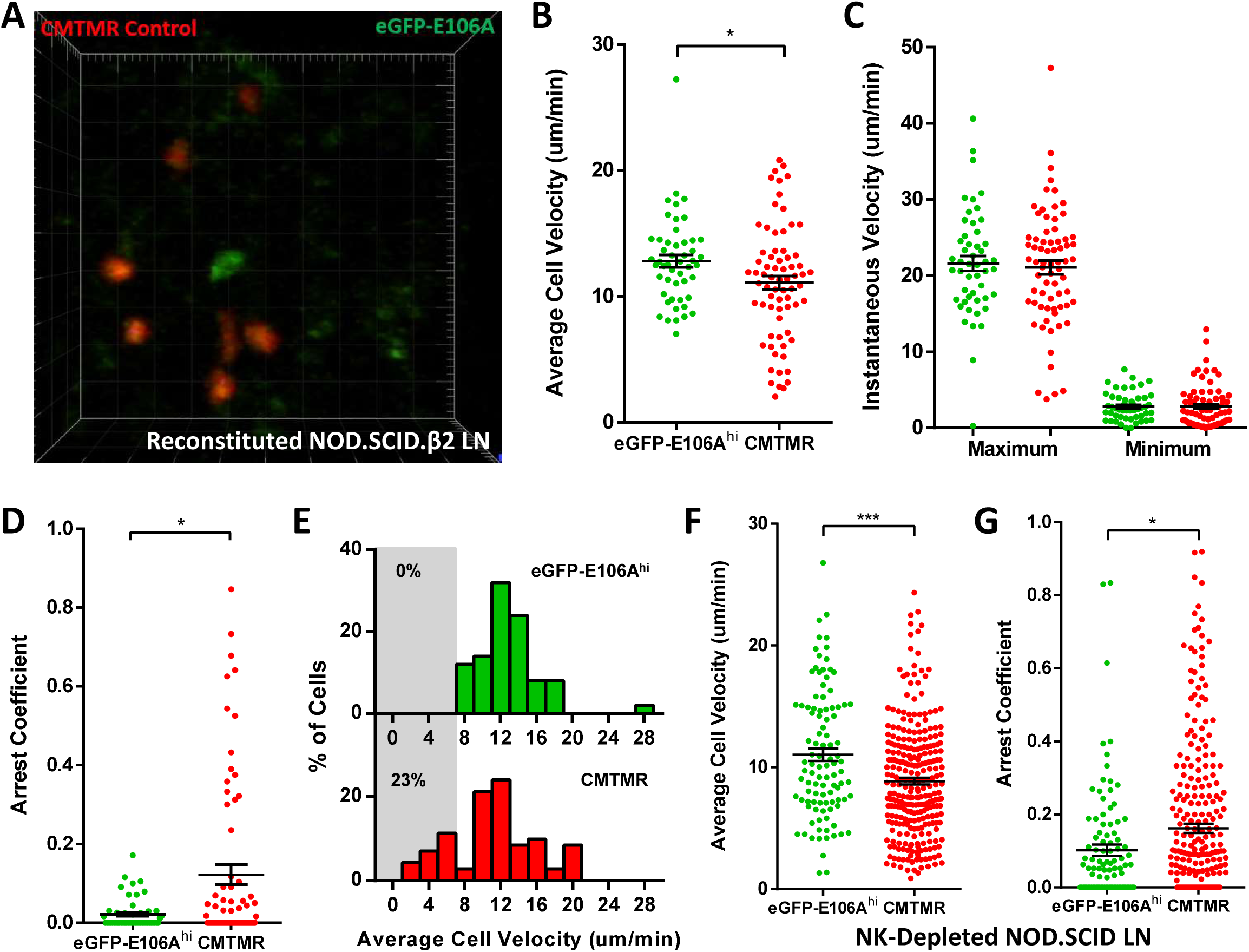
Orai1 block increases human T cell motility within reconstituted NOD.SCID.β2 lymph nodes. (**A**) Two-photon microscopy of migrating human T cells, showing eGFP-E106A transfected cells in green and CMTMR-labeled mock transfected cells in red, within intact mouse lymph node 18 hours after adoptive co-transfer of 5×10^6^ of each cell type. (**B**) Average cell velocities of eGFP-E106A^hi^ (n = 50) versus CMTMR-labeled control (n = 71) T cells; bars represent mean ± SEM, data from independent experiments using 4 different donors. (**C**) Maximum and minimum cellular instantaneous velocities of eGFP-E106A^hi^ (green) versus CMTMR-labeled (red) control T cells. (**D**) Arrest coefficients of eGFP-E106A^hi^ compared with CMTMR-labeled control T cells, defined as fraction of time with instantaneous velocity < 2 μm/min. (**E**) Frequency distribution of average cell velocities for eGFP-E106A^hi^ (top) and CMTMR-labeled control T cells (bottom), cells with average velocity < 7 μm/min are highlighted in gray; tick marks denote the center of every other bin. (**F,G**) Average cell velocities (**F**) and arrest coefficients (**G**) of eGFP-E106A^hi^ (green, n = 102) versus CMTMR-labeled control (red, n = 278) human T cells in NK cell depleted immunodeficient mouse lymph nodes; bars represent mean ± SEM, data from independent experiments using 8 different donors, *** = p < 0.005.

To replicate our findings in a different immunodeficient mouse model, we repeated our human T cell adoptive transfer protocol using NOD.SCID mice depleted of NK cells. Lymph nodes in these mice are small and contain reticular structures but are completely devoid of lymphocytes (Shultz, Schweitzer et al. 1995). Similar to experiments on reconstituted NOD.SCID.β2 mice, eGFP-E106A^hi^ human T cells in NOD.SCID lymph nodes migrated with elevated average velocities compared to control T cells (11.0 ± 0.5 μm/min vs. 8.8 ± 0.3 μm/min, p = 0.0004; Figure 2F), and exhibited lower arrest coefficients (0.10 ± 0.02 vs. 0.16 ± 0.01, p = 0.0516; Figure 2G). Both eGFP-E106A^hi^ and control T cells migrated at lower speeds in the NK-depleted NOD.SCID model compared to the reconstituted NOD.SCID.β2 model. Because control human T cells in reconstituted NOD.SCID.β2 lymph nodes migrated at similar speeds to what we have previously seen for wildtype mouse T cells *in vivo* (Miller, Wei et al. 2002), reconstitution results in a lymph node environment that more closely mimics normal physiological conditions. Furthermore, the greater effect of Orai1 block on T cell arrest coefficients in crowded reconstituted lymph nodes suggests that Orai1’s role in motility is more pronounced in crowded cell environments.

### Orai1 channel activity triggers pauses during human T cell motility *in vitro* in the absence of extrinsic cell contact

To evaluate whether the pronounced effect of Orai1 channel block on the arrest coefficient in reconstituted lymph nodes was a result of environmental factors such as increased cellular contacts or increased confinement, we tracked human T cells in microfabricated PDMS chambers with cell-sized microchannels 7 µm high × 8 µm wide. These ICAM-1 coated microchannels simulate the confined environment of densely packed lymph nodes (Jacobelli, Friedman et al. 2010), while eliminating possible cell-extrinsic factors. Transfected human T cells were activated with plate-bound anti-CD3/28 antibodies and soluble IL-2, then dropped into chambers and monitored by time-lapse confocal microscopy, using phase contrast to visualize eGFP-E106A^lo^ T cells (Figure 3A,B). Upon entry into microchannels, eGFP-E106A^hi^ T cells migrated with higher average cell velocities than eGFP-E106A^lo^ T cells (14.2 ± 0.6 μm/min vs. 10.9 ± 0.5 μm/min, p < 0.0001; Figure 3C), similar to our *in vivo* findings from intact lymph node (*c.f*., Figures 3C, 2B). To ensure that the observed difference in cell velocity was due to suppressed Orai1 channel function and not overexpression of Orai1 protein, we also tracked T cells transfected with eGFP-tagged wildtype Orai1. Both eGFP-Orai1^hi^ and eGFP-Orai1^lo^ T cells migrated at the same average cell velocity (10.7 ± 0.8 μm/min vs. 10.5 ± 0.8 µm/min; Hodges-Lehmann median difference of -0.84 µm/min, -2.96 to 1.28 µm/min 95%CI; Figure 3C), demonstrating that Orai1 channel overexpression, in itself, does not perturb T cell motility in microchannels. Since eGFP-E106A^lo^ T cells have reduced Orai1 channel activity but still retain the same cell velocity as eGFP-Orai1 transfected T cells (*c.f*., Figures 1A, 3C), this suggests that partial Orai1 function is sufficient to generate normal pausing frequency in confined environments. The frequency distribution of cell velocities *in vitro* is comparable to our *in viv*o data: fewer GFP-E106A^hi^ T cells migrated with average cell velocities < 7 µm/min as compared to eGFP-E106A^lo^ T cells (11% vs 29%; *c.f*., Figures 3D, 2E). Furthermore, eGFP-E106A^hi^ T cells exhibited lower arrest coefficients (0.05 ± 0.01 vs. 0.08 ± 0.01, p = 0.0015; Figure 3E) and less variation in velocity than eGFP-E106A^lo^ T cells (39.5 ± 1.9 % vs. 45.1 ± 1.6 %, p = 0.0138; Figure 3F). Although eGFP-E106A^hi^ T cells had lower arrest coefficients, the durations of their pauses were not significantly different than in eGFP-E106A^lo^ T cells (Hodges-Lehmann median difference of 0 seconds, -8.43 to 4.71 seconds 95%CI; Figure 3G). Taken together, the reduced arrest coefficients in eGFP-E106A^hi^ T cells without any change in the durations of pauses indicate that inhibition of Orai1 channel activity results in reduced frequency of pauses during T cell motility. These *in vitro* results confirm our *in vivo* findings and support the hypothesis that Orai1 activity intermittently triggers cell arrest, resulting in an overall decrease in motility within confined environments. Moreover, since our *in vitro* microchannel assay eliminates extrinsic cell-cell interactions, this indicates that Orai1 is activated in a cell-intrinsic manner when modulating T cell motility.

**Figure 3.**
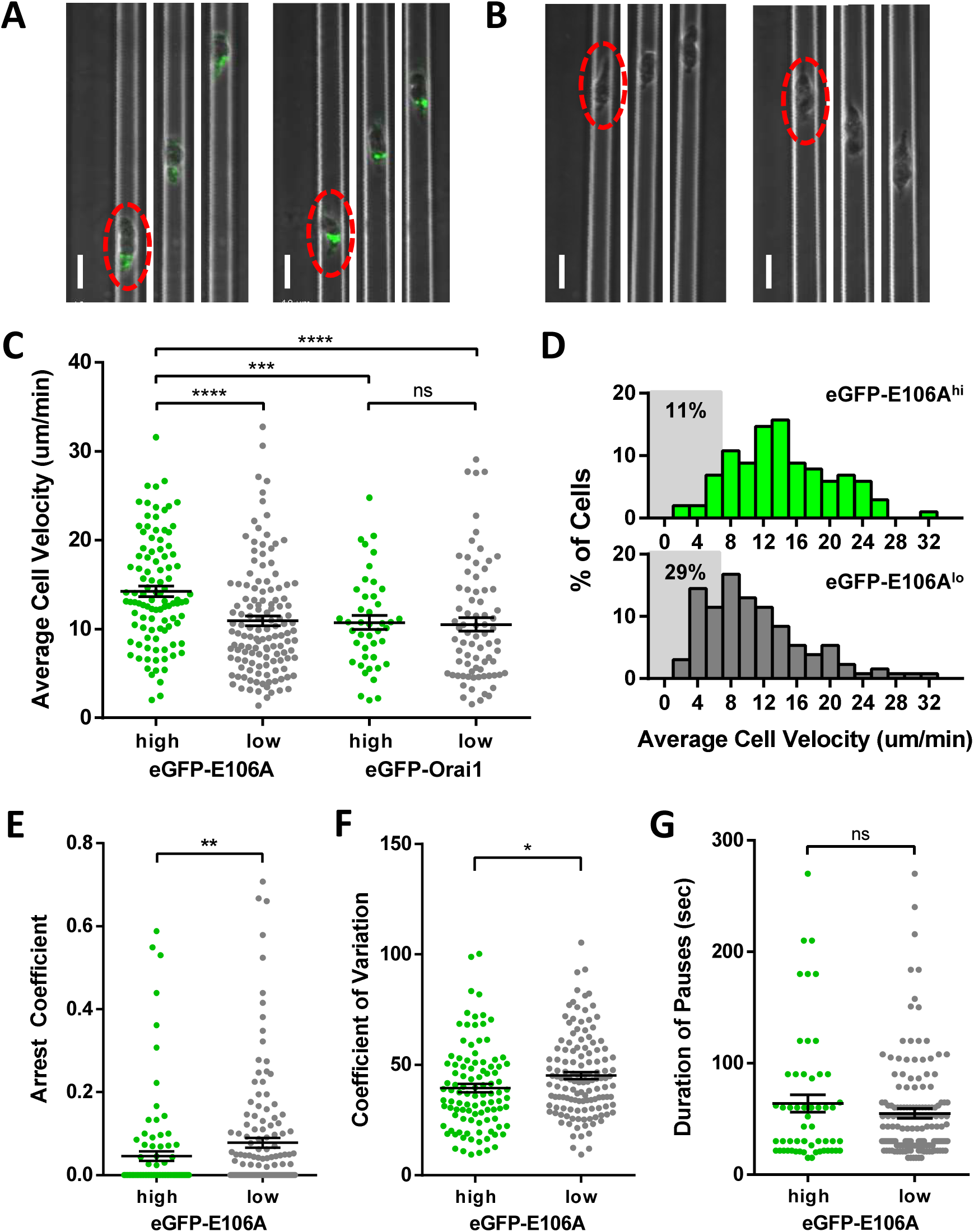
Orai1 block reduces frequency of pausing during human T cell motility *in vitro*. (**A,B**) Confocal microscopy of eGFP-E106A transfected human CD4^+^ T cells in microfabricated channels 7 μm high by 8 μm wide, showing two individual eGFP-E106A^hi^ T cells (**A**) and two eGFP-E106A^lo^ T cells (**B**), each circled in red in the first frame; individual images taken 1 min apart, scale bar = 10 μm. (**C**) Comparison of average cell velocities of eGFP-E106A transfected T cells (eGFP-E106A^hi^ cells in green, n = 102; eGFP-E106A^lo^ cells in gray, n =131) vs eGFP-Orai1 transfected control T cells (eGFP-Orai1^hi^ cells in green, n = 43; eGFP-Orai1^lo^ cells in gray, n = 76); bars represent mean ± SEM, data from independent experiments using 5 different donors. (**D**) Frequency distribution of average cell velocities of eGFP-E106A^hi^ (top) and eGFP-E106A^lo^ (bottom) human T cells, cells with average velocity < 7 μm/min are highlighted in gray; tick marks denote the center of every other bin. (**E**) Arrest coefficients of eGFP-E106A^hi^ vs eGFP-E106A^lo^ human T cells, defined as fraction of time each individual cell had an instantaneous velocity < 2 μm/min. (**F**) Variance in velocity of eGFP-E106A^hi^ vs eGFP-E106A^lo^ human T cells, coefficient of variation is calculated by standard deviation divided by the mean of instantaneous velocity for each individual cell. (**G**) Duration of pauses for eGFP-E106A^hi^ vs eGFP-E106A^lo^ human T cells; bars represent mean ± SEM, * = p < 0.05, ** = p < 0.01, *** = p < 0.005, **** = p < 0.001.

### A novel ratiometric genetically encoded Ca^2+^ indicator, Salsa6f

In order to develop a better tool to monitor Ca^2+^ signaling in T cells both *in vivo* and *in vitro*, we took advantage of the latest generation of genetically encoded Ca^2+^ indicators (GECIs) (Zhao, Araki et al. 2011, Chen, Wardill et al. 2013). A variety of single fluorescent protein-based GECIs were transiently expressed and screened in HEK 293A cells (Figure 4A), and GCaMP6f was selected based on fluorescence intensity, dynamic range, and Ca^2+^ affinity (*k*_*d*_ = 373 nM) suitable for detecting a variety of cytosolic Ca^2+^ signals. To enable cells to be tracked even when their basal Ca^2+^ level evokes little GCaMP6f fluorescence, we fused GCaMP6f to the Ca^2+^-insensitive red fluorescent protein tdTomato, chosen for its photostability and efficient two-photon excitation (Drobizhev, Makarov et al. 2011). A V5 epitope tag (Lobbestael E 2010) serves to link tdTomato to GCaMP6f (Figure 4C). The resultant ratiometric fusion indicator, coined “Salsa6f” for the combination of red tdTomato with the green GCaMP6f, was readily expressed by transfection into HEK 293A cells or human T cells. Salsa6f exhibited a ten-fold dynamic range, with a brightness comparable to GCaMP6f alone (Figure 4A,B). For two-photon microscopy, both components of Salsa6f can be visualized by femtosecond excitation at 900 nm (Figure 4D). GCaMP6f produces increased green fluorescence during elevations in cytosolic Ca^2+^, while tdTomato provides a stable red fluorescence that facilitates cell tracking and allows for ratiometric Ca^2+^ imaging (Figure 4D, **Video 1**). Salsa6f is excluded from the nucleus, ensuring accurate measurement of cytosolic Ca^2+^ fluctuations (Figure 4D). When expressed in human T cells, Salsa6f reported Ca^2+^ oscillations induced by immobilized anti-CD3/28 antibodies with a high signal to noise ratio and time resolution (Figure 4E,F).

**Figure 4.**
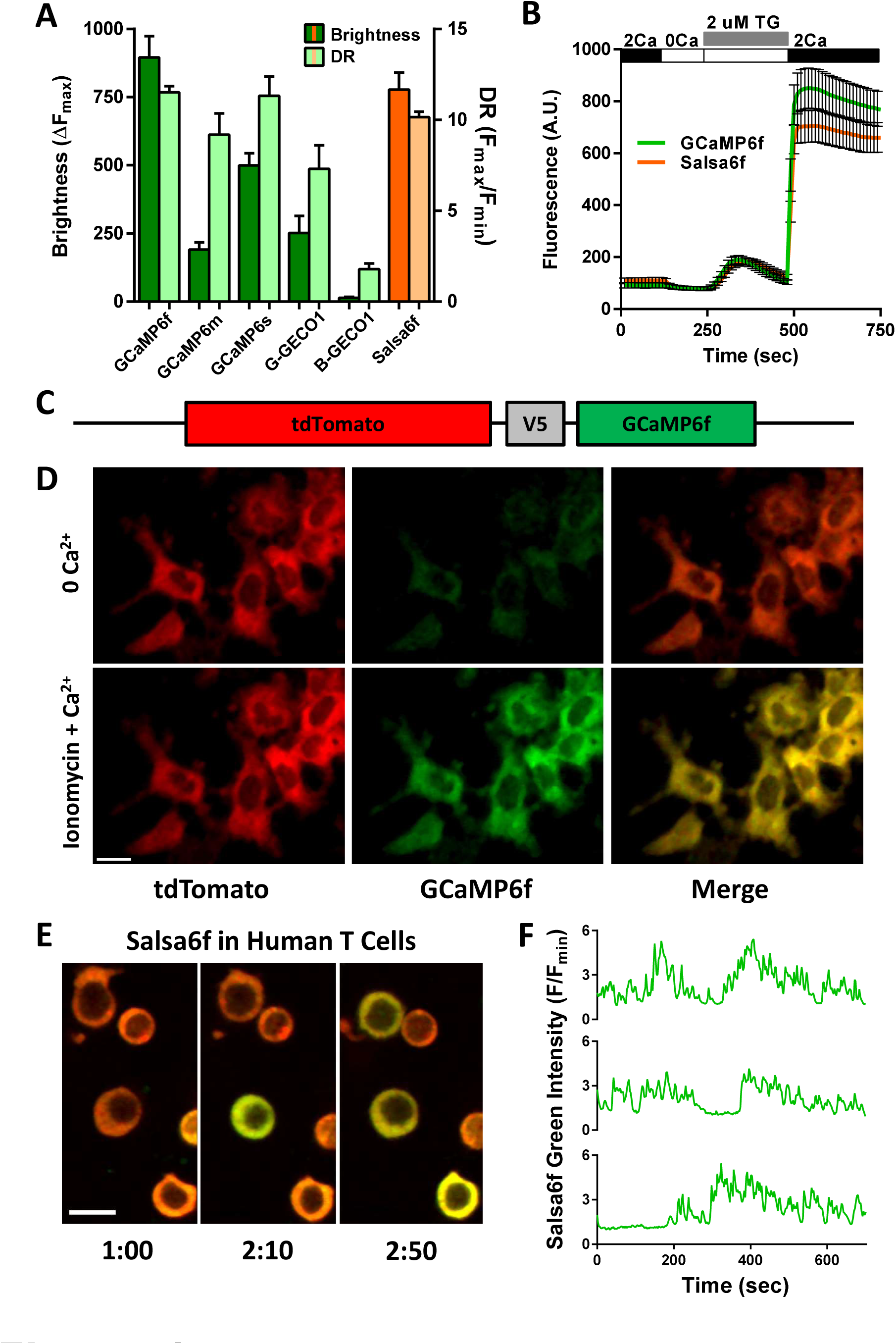
Design of novel tdTomato-V5-GCaMP6f fusion probe “Salsa6f” and characterization in living cells. (**A**) Several genetically encoded Ca^2+^ indicators were screened *in vitro* in HEK 293A cells, by co-transfecting with Orai1/STIM1 and measuring Ca^2+^ influx after thapsigargin-induced store depletion, showing maximum change in fluorescence intensity in dark green bars and dynamic range (DR) in light green bars, with Salsa6f shown in orange bars on right; n > 30 cells per probe, from two different transfections, error bars indicate SEM. (**B**) Averaged thapsigargin-induced Ca^2+^ entry, measured by change in GFP fluorescence, in GCaMP6f (green, 11.52 ± 0.34, n = 63) or Salsa6f (orange, 10.16 ± 0.31, n = 78) transfected HEK cells; data from two different transfections, error bars indicate SEM. (**C**) Diagram of Salsa6f construct used in transfection. (**D**) Two-photon images of Salsa6f co-transfected in HEK cells with Orai1/STIM1, showing red (tdTomato), green (GCaMP6f), and merged channels, at baseline in 0 mM extracellular Ca^2+^ and after maximum stimulation with 2 mM ionomycin in 2 μM extracellular Ca^2+^; scale bar = 20 μm; see **Video 1**; data representative of at least three different experiments. (**E**) Confocal time lapse microscopy of human CD4^+^ T cells transfected with Salsa6f, then activated for two days on platebound anti-CD3/28 antibodies; time = min:sec, scale bar = 10 μm. (**F**) Representative cell traces of activated Salsa6f transfected human T cells, tracking green fluorescence intensity; data representative of at least three different experiments.

### Spontaneous Ca^2+^ signals during confined motility *in vitro* are correlated with reduced T cell velocity

Human CD4^+^ T cells were transfected with Salsa6f, activated for two days with plate-bound anti-CD3/28 antibodies, then dropped into ICAM-1 coated microchambers. As previously shown (Figure 4E), Salsa6f is localized to the cytosol, with red fluorescence from tdTomato that reflects fluctuations in cell movement and very low baseline green fluorescence from GCaMP6f that rises sharply during Ca^2+^ signals (Figure 5A-D). Salsa6f-transfected human T cells were tracked in both confined microchannels (Figure 5A, **Video 2**) and the open space adjacent to entry into microchannels (Figure 5C, **Video 3**), to evaluate T cell motility under varying degrees of confinement. Intracellular Ca^2+^ levels were monitored using the ratio of total GCaMP6f fluorescence intensity over total tdTomato fluorescence intensity, enabling detection of a notably stable baseline ratio unaffected by motility artifacts in moving T cells while reporting spontaneous Ca^2+^ signals that could be compared to changes in motility (Figure 5B,D, orange and black traces, respectively).

**Figure 5.**
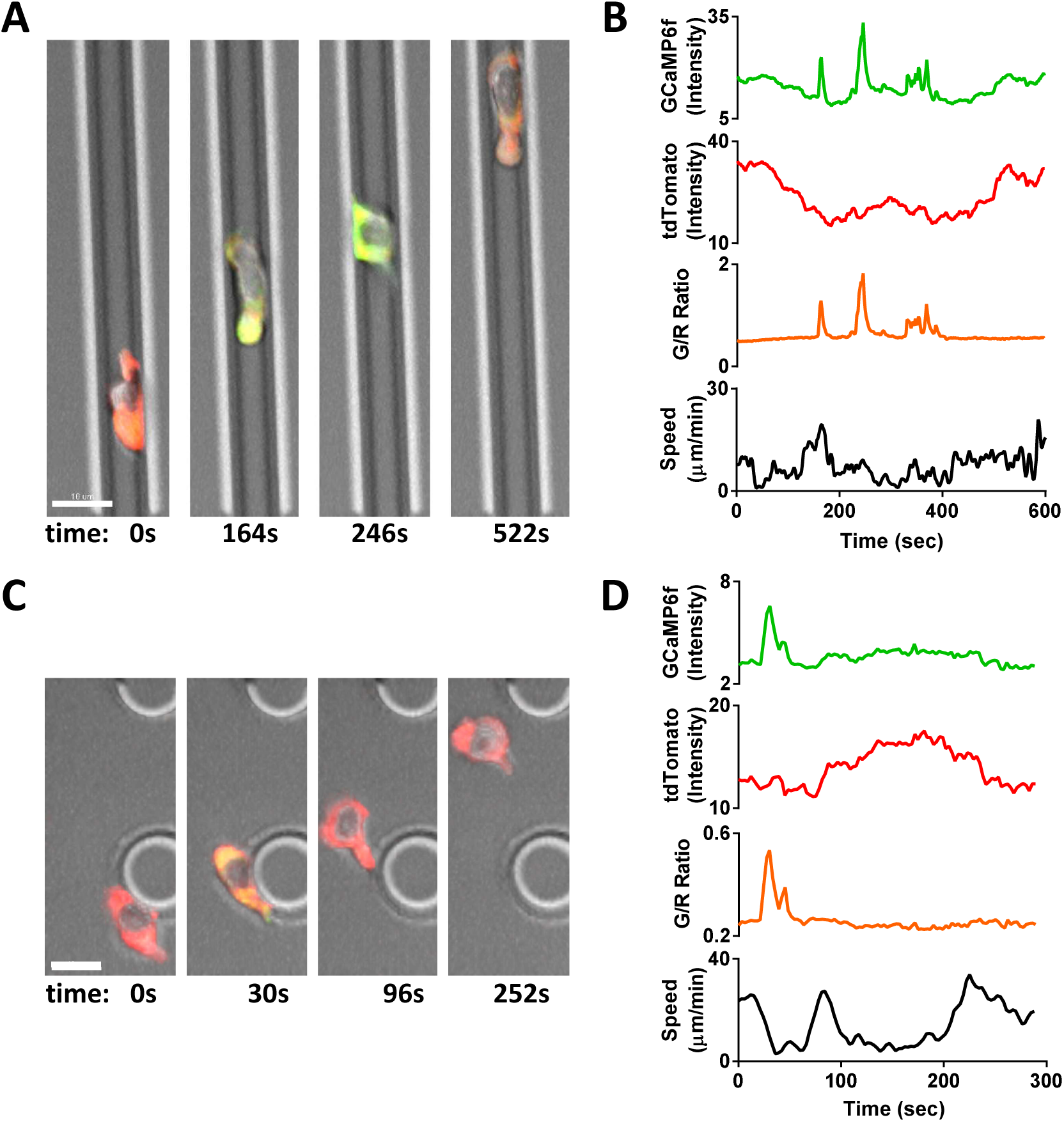
Tracking Ca^2+^ signals in human T cells *in vitro* with Salsa6f. (**A,C**) Confocal microscopy of Salsa6f transfected human CD4^+^ T cells in ICAM-1 coated microchannels 7 μm high by 8 μm wide (**A**) and open space (**C**), showing merged red (tdTomato), green (GCaMP6f), and DIC channels; circular structures shown in (**C**) are support pillars part of the PDMS chamber; scale bar = 10 μm, time = sec; see **Videos 2 and 3**. (**B,D**) Total intensity tracings of GCaMP6f (green) and tdTomato (red) fluorescence, GCaMP6f/tdTomato ratio (orange), and speed (black), for corresponding T cells shown in (**A**) and (**C**); data representative of independent experiments from three different donors.

Human T cells expressing Salsa6f migrating in confined microchannels exhibited stable baseline cytosolic Ca^2+^ levels (Figure 6A), although sporadic Ca^2+^ signals were apparent as brief peaks unrelated to changes in motility (Figure 6B, arrowheads), or as more sustained periods of Ca^2+^ elevation associated with reduced cell velocity (Figure 6B, gray highlights). To evaluate the correspondence between T cell velocity and Ca^2+^ signals, we compared average T cell velocities during periods of sustained Ca^2+^ elevations to average velocities at baseline Ca^2+^ levels. T cell velocity decreased significantly when cytosolic Ca^2+^ was elevated above baseline (5.9 ± 0.1 μm/min vs. 10.0 ± 0.1 μm/min, p < 0.0001; Figure 6C). Ca^2+^ signaling episodes that last for 30 seconds or longer accompany and appear to closely track the duration of pauses in cell movement. Comparison of instantaneous velocities with corresponding cytosolic Ca^2+^ signals by scatter plot revealed a strong inverse relationship: highly motile T cells always exhibited baseline Ca^2+^ levels, while elevated Ca^2+^ levels were only found in slower or arrested T cells (Figure 6D). It is important to note that these Ca^2+^ signals and reductions in velocity occurred in the absence of any extrinsic cell contact or antigen recognition, indicating that Ca^2+^ elevations, like pausing and Orai1 activation, are triggered in a cell-intrinsic manner.

**Figure 6.**
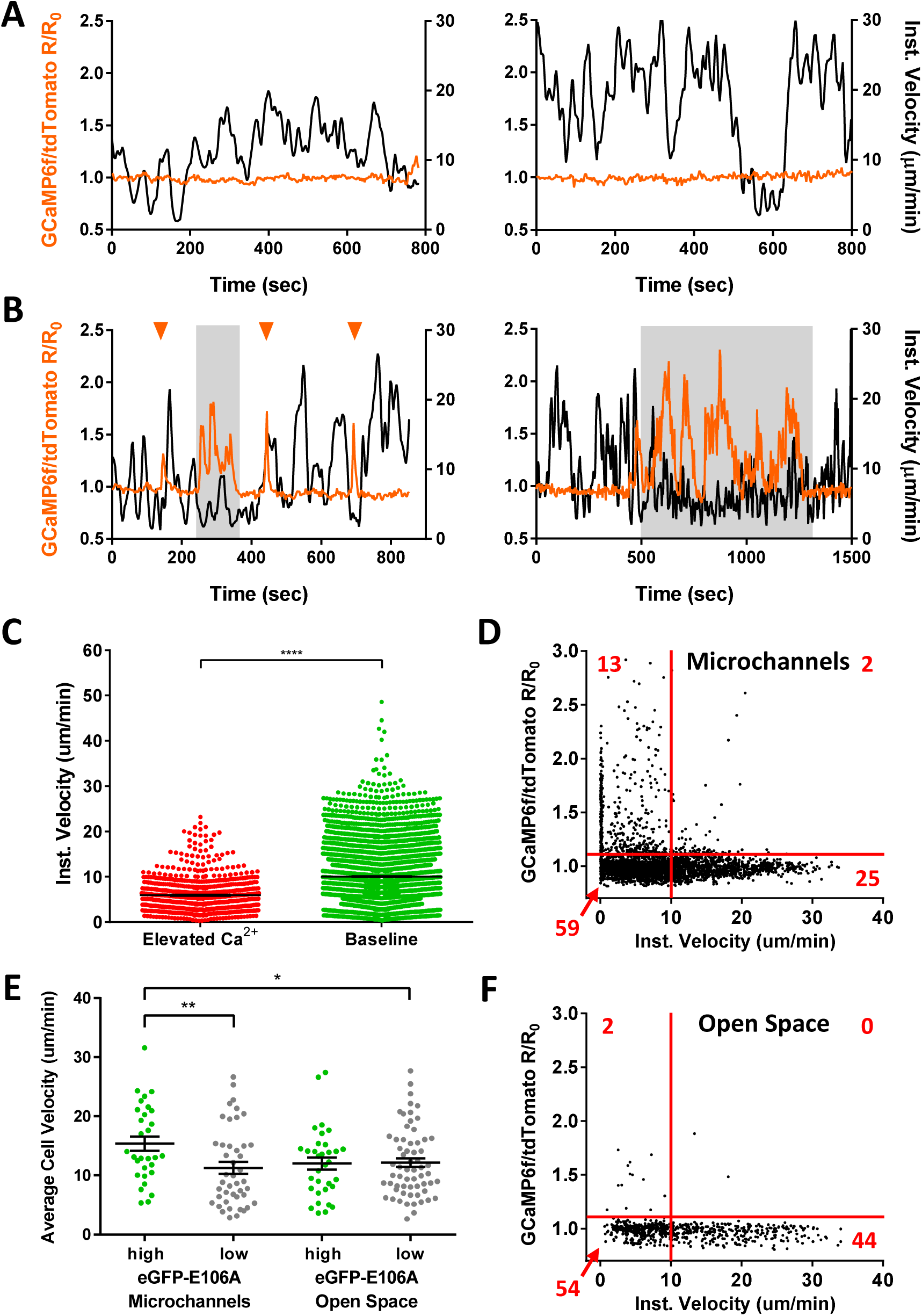
Spontaneous Ca^2+^ signals during human T cell motility *in vitro* are correlated with reduced velocity. (**A,B**) Sample tracks from Salsa6f transfected human T cells in microchannels, with intracellular Ca^2+^ levels measured by R/R_0_ of GCaMP6f over tdTomato fluorescence intensity (orange), overlaid with instantaneous cell velocity (black), cells in (**A**) have stable Ca^2+^ levels, cells in (**B**) show brief Ca^2+^ transients (arrowheads) or sustained Ca^2+^ signaling (gray highlights). (**C**) Instantaneous velocity of Salsa6f transfected human T cells in microchannels during elevated cytosolic Ca^2+^ levels (red) and during basal Ca^2+^ levels (green); n = 22 cells, data from independent experiments using three different donors; **** = p < 0.001. (**D**) Scatter plot of Salsa transfected human T cells in microchannels, instantaneous cell velocity versus GCaMP6f/tdTomato R/R_0_ for each individual time point analyzed; red numbers in each quadrant show percent of time points, split by 1.10 R/R_0_ and 10 μm/min; n = 4081 points. (**E**) Mean track velocity of eGFP-E106A transfected human T cells, comparing eGFP-E106A^hi^ (green) versus eGFP-E106A^lo^ T cells (gray) in confined microchannels vs open space; n = 30, 44, 33, and 62 cells, respectively; bars represent mean ± SEM, data from independent experiments using two different donors, * = p < 0.05, ** = p < 0.01. (**F**) Scatter plot of Salsa transfected human T cells in open space, instantaneous cell velocity versus GCaMP6f/tdTomato R/R_0_ for each individual time point analyzed; red numbers in each quadrant show percent of cells, split by 1.10 R/R_0_ and 10 μm/min; n = 723 points.

To compare the effects of Orai1 activity on the motility of T cells in different *in vitro* environments, we also monitored T cell migration within the open space in PDMS chambers adjacent to entry into microchannels (*c.f*., Figure 5A,C). We reasoned that in this two-dimensional space with reduced confinement, T cells may not gain sufficient traction for rapid motility, and instead may favor integrin-dependent sliding due to increased exposure to the ICAM-1 coated surface (Krummel, Friedman et al. 2014). In addition, the same population of T cells could be tracked as they migrated into and along the confined microchannels, providing a valuable internal control. We found that eGFP-E106A^hi^ T cells migrated with similar velocities to eGFP-E106A^lo^ T cells in the open space (12.0 ± 1.0 μm/min vs. 12.2 ± 0.7 μm/min; Hodges-Lehmann median difference of 0.15 μm/min, -2.46 to 2.40 μm/min 95%CI; Figure 6E), but these eGFP-E106A^hi^ T cells still exhibited higher motility in the microchannels than eGFP-E106A^lo^ T cells (15.4 ± 1.2 μm/min vs. 11.3 ± 1.0 μm/min; p = 0.0099; Figure 6E). Furthermore, Salsa6f transfected T cells within the open space rarely produced Ca^2+^ transients that coincided with reduced instantaneous velocity (*c.f*., Figure 6D,F, top left quadrants, 13% of the time in microchannels vs 2% in open space), implying that Ca^2+^ elevations, and by extension, Orai1 channel activity, do not generate pauses when T cells are reliant on integrin binding for motility. Taken together, these experiments establish a role for the cell-intrinsic activation of Orai1 channels in modulating T cell motility within confined environments.

## Discussion

In this study, we demonstrate that Orai1 channel activity regulates motility patterns that underlie immune surveillance, specifically within confined environments. Human T cells expressing the dominant-negative Orai1-E106A construct migrated with higher average velocities than controls, both *in vivo* in immunodeficient mouse lymph nodes and *in vitro* in confined microchannels. In particular, we found that the increase in average cell velocity was not due to an increase in maximum cell velocity, but to a reduced frequency of cell arrest. Furthermore, we introduce a novel ratiometric genetically encoded Ca^2+^ indicator, Salsa6f, and show in confined microchannels devoid of cell-extrinsic factors, that human T cells exhibit spontaneous Ca^2+^ signals that coincide with reduced cell velocity. We propose that cell-intrinsic activation of Orai1 channel activity regulates the timing of stop-and-go motility in T cells and tunes the search for cognate antigen in the crowded lymph node.

Salsa6f preserves the exceptional performance of GCaMP6f, with similar brightness and dynamic range, and adds to it the bright fluorescence of tdTomato, allowing cell tracking in the absence or presence of Ca^2+^ signaling. FRET based ratiometric indicators (Heim and Griesbeck 2004, Thestrup, Litzlbauer et al. 2014) lack the brightness and dynamic range of Salsa6f. Exclusion of Salsa6f from the nucleus enables precise monitoring of cytosolic Ca^2+^ signals. In addition, we observed no overt effects of Salsa6f expression on cell behavior. Ratiometric imaging of Salsa6f eliminates motility artifacts and reveals stable baseline levels of Ca^2+^ in the cytoplasm of rapidly moving human T cells; this finding enables Ca^2+^ signals to be identified clearly and their temporal progression followed precisely. Moreover, both GCaMP6f and tdTomato components of Salsa6f can be visualized in two-photon microscopy by femtosecond excitation at 900nm. The genetically encoded nature of Salsa6f allows it to be continuously expressed in stably integrated cell lines or transgenic reporter animals, enabling tracking of cell fates and long term studies. Taken together, these considerations indicate that Salsa6f is well suited for assessing links between Ca^2+^ signaling and T cell motility and activation, and demonstrate that Salsa6f could be usefully employed to study Ca^2+^ signaling dynamics in many other cell types.

Our Orai1-E106A construct derives its specificity and potency by targeting the pore residue responsible for the channel selectivity filter. Incorporation of Orai1-E106A likely blocks heterodimers of Orai1 and other channels, such as Orai2 or Orai3, in addition to homomeric Orai1 channels. Very recent evidence demonstrates the existence of heteromeric channels composed of Orai1 and Orai2 in T cells (Vaeth, Yang et al. 2017). These heteromers appear to simply reduce the flow of Ca^2+^ through the Orai1 channel without targeting additional signaling pathways. In the absence of contradictory evidence, we conclude that in T cells Orai1-E106A acts to produce an essentially complete functional knockdown of Orai1-mediated store-operated Ca^2+^ entry.

In lymph nodes, T cells experience a dense and crowded environment packed with fibroblastic reticular cells, resident dendritic cells, and numerous other migrating immune cells (Miller, Wei et al. 2002, Bajenoff, Egen et al. 2006). Over the last decade there has been a developing realization that mechanisms of motility in the lymph node differ in important ways from mechanisms used in less densely packed tissues and in standard, adhesion-rich *in vitro* assays. First, integrins – key determinants of lymphocyte adhesion and key components of *in vitro* motility assays – are dispensable for T cell and dendritic cell motility in the lymph node, where the absence of shear forces prevent stable integrin adhesiveness (Woolf, Grigorova et al. 2007, Lammermann, Bader et al. 2008). Second, theoretical and experimental work suggested the existence of a novel motility mode in which confined cells brace against their neighbors to push through crowded environments (Hawkins, Piel et al. 2009). Third, development of an *in vitro* assay using microchannels has enabled cell confinement to be varied *in vitro*, allowing confinement to be studied in isolated cells and mimicking the effects of cell crowding inside intact tissues such as the lymph node (Jacobelli, Friedman et al. 2010). Under confined conditions that elicit maximal cell velocities, T cells utilize high speed “amoeboid walking” which relies on multiple brief contacts to substrate, independent of integrin function. This motility mode appears to be driven by actin-network expansion and contraction of myosin IIA motors, which facilitates protrusive flowing of the cell’s leading edge (Jacobelli, Friedman et al. 2010, Krummel, Friedman et al. 2014). Taken together, these studies indicate that amoeboid walking is the predominant mode of T cell motility within lymph nodes.

Alternatively, when abundant extracellular adhesion molecules are available, T cells favor an integrin-dependent motility mode termed “haptokinetic sliding”, such as in peripheral tissues, or in open field cell motility assays (Svensson, McDowall et al. 2010, Krummel, Friedman et al. 2014). Haptokinetic sliding is generally slower than amoeboid walking and relies upon a single continuous zone of contact to substrate. Integrin-dependent open field *in vitro* assays have been used to highlight the accumulation of K_Ca_3.1 and TRPM7 channels at the uropod where the Ca^2+^ dependent protease calpain-2 may regulate turnover of LFA-1 adhesions (Svensson, McDowall et al. 2010, Kuras, Yun et al. 2012). While these integrin-dependent assays reportedly exclude a role for Orai1 in T cell motility, we argue that they do not reflect the predominant mode of motility used *in vivo* in the lymph node, and so do not bear directly upon the involvement of Orai1 in T cell motility in the lymph node. However, assessing integrin-dependent motility in parallel with confinement-dependent motility allows identification of motility mode-specific mechanisms and controls for cell health and cytoskeletal integrity.

Many studies demonstrate that Ca^2+^ influx through Orai1 is a signal that selectively activates downstream effectors, including calmodulin, calcineurin, and the transcription factor NFAT (Dolmetsch and Lewis 1994, Negulescu, Shastri et al. 1994, Dolmetsch, Lewis et al. 1997, Kar, Samanta et al. 2014). We observe inhibition of pausing by Orai1 block only under confined conditions both *in vitro* and *in vivo*; this indicates that Orai1 block is not generally deleterious for cell health and movement, but instead acts upon subcellular mechanisms that are selectively employed during confined motility. The maximum cell velocity and mean pause length are unchanged by Orai1 block. Instead, Orai1 block selectively alters the timing of when pauses are triggered during T cell motility. The selective alteration of timing by mutation is a hallmark of cell regulatory pathways and, together with the widespread signaling roles already ascribed to Orai1 in other systems, establishes that Orai1 channel activity acts to regulate T cell motility. This model is supported by our Ca^2+^ imaging data: episodes of Ca^2+^ signaling lasting longer than 30 s coincide with pauses in cell movement. In many contexts, Ca^2+^ signaling has been to shown not only accompany, but also to cause, cell arrest and loss of cell polarity, such as in T cells after activation by antigen (Negulescu, Krasieva et al. 1996, Dustin, Bromley et al. 1997, Wei SH 2007). While we do not show that the Ca^2+^ signals we observe emanate directly from Orai1 channels, taken together our data are consistent with Orai1 actively regulating cell motility by directly inducing subcellular mechanisms that lead to cell arrest.

In previous studies of Orai1 signaling, Orai1 activation has been placed downstream of extracellular ligand binding to cell surface receptors, integrating their input upon use-dependent depletion of Ca^2+^ from the ER (Feske 2007, Cahalan and Chandy 2009). Using *in vitro* microchannel assays, we find that Orai1 block alters motility in the absence of ongoing cell-cell interactions. Therefore, Orai1 is activated in an unknown cell-intrinsic manner; at this point it is unclear which aspects of internal cell state lead to Orai1 channel opening and pauses in motility. Our identification of Orai1 provides a clear new entry into mechanisms that drive T cells motility patterns used for immune surveillance. Of particular interest is determining the step in the signaling cascade from phospholipase C to Orai1 that is targeted by this novel activation pathway.

The apparently random nature of naïve T cell movement in the lymph node has led to the hypothesis that T cells use intrinsic and stochastic motility mechanisms to accomplish immune surveillance (Wei, Parker et al. 2003, Mrass, Petravic et al. 2010, Germain, Robey et al. 2012). Because of recent studies, our understanding of the molecular mechanisms that produce intrinsic motility patterns is beginning to take shape (Jacobelli, Friedman et al. 2010, Gerard, Patino-Lopez et al. 2014). Myosin 1g, like Orai1, is selectively required for cell-intrinsic motility mechanisms under confined conditions (Gerard, Patino-Lopez et al. 2014). Yet differences in phenotype between Orai1 and Myo1g block suggest that these proteins act in different, and in part opposing, ways to control T cell motility. While Orai1 block reduces pausing but does not alter T cell velocity, Myo1g block increases pausing and causes cells to move faster. Our observations support the assertion that Myo1g and Orai1 act in parallel cell-intrinsic pathways that jointly modulate T cell motility.

Our identification of an intrinsic Orai1-dependent T cell motility program provides further support for the hypothesis that intrinsic and stochastic T cell motility patterns underlie immune surveillance in the lymph node. Immune surveillance requires balancing many factors associated with antigen search, including speed and sensitivity. Myo1g activity biases surveillance toward sensitivity by increasing the duration of individual T cell-dendritic cell contacts. Pauses in motility caused by Orai1 activity could allow similar biases in surveillance by increasing the fraction of time that, collectively, T cells are stationary and available for extended contacts with dendritic cells. Moreover, in the absence of other motility changes, the interval between Orai1-dependent pauses determines the distance T cells travel between pauses. In this way Orai1 channel activity could tailor T cell excursions to match the density and reach of dendritic cells in the lymph node.

## Acknowledgments

We thank Drs. Luette Forrest and Olga Safrina for expert assistance, excellent animal care and vivarium support, and Dr. Audrey Gerard, the Matthew Krummel Lab at UCSF, and the Christopher Hughes Lab at UCI for assistance in establishing the microchamber fabrication technique. We acknowledge the UC Irvine Institute for Clinical and Translational Science, and Dr. Jennifer Atwood of the Flow Core Facility supported by the UC Irvine Institute of Immunology.

## Methods

### Human T cell purification and transfection

Human PBMCs were isolated from blood of voluntary healthy donors by Histopaque-1077 (1.077 g/mL; Sigma, St. Louis, MO) density gradient centrifugation. Human CD3^+^, CD4^+^, or CD8^+^ T cells were isolated using the appropriate EasySep Human T Cell Isolation Kit (StemCell Technologies, Vancouver, Canada) according to manufacturer’s instructions. The purity of isolated cells was confirmed to be >95% by flow cytometry. Purified cells were rested overnight in complete RPMI, then transfected by nucleofection (Lonza, Walkersville, MD), using the high-viability “U-014” protocol. Enhanced green fluorescent protein (eGFP)-tagged wildtype Orai1, eGFP-tagged Orai1-E106A mutant, Salsa6f (tdTomato-V5-GCaMP6f construct), or empty vector control were transfected as indicated. Human T cells were used for experiments 3-48 hr after transfection. For *in vitro* imaging experiments, T cells were rested for 3-4 hr in complete RPMI as indicated in the manufacturer’s instructions, then washed and activated on plate-bound αCD3 and αCD28 (Tonbo Biosciences, San Diego, CA) in 2.5 ng/mL recombinant human IL-2 (BioLegend, San Diego, CA), and imaged 24-48 hr after transfection.

### Mice

NOD.Cg-*Prkdc^scid^B2m^tm1Unc^*/J (NOD.SCID.β2) and NOD.CB17-*Prkdc^scid^*/J (NOD.SCID) mice obtained from Jackson Laboratory (Stock #002570 and #001303) were housed and monitored in a selective pathogen-free environment with sterile food and water in the animal housing facility at the University of California, Irvine. NOD.SCID.β2 mice were reconstituted with human peripheral blood leukocytes (PBLs) as described previously (Mosier, Gulizia et al. 1988). A total of 3×10^7^ human PBLs were injected i.p., and experiments were performed three weeks later. To inhibit NK cell activity, NOD.SCID mice were i.p. injected with 20 μL anti-NK cell antibody (rabbit anti-Asialo GM1, Wako Chemicals, Irvine, CA) according to manufacturer’s instructions 3-4 days before adoptive transfer of human T cells. Mice used were between 8 and 18 weeks of age. For adoptive transfer experiments, 5×10^6^ human CD3^+^ T cells were labeled with 10 μΜ CellTracker CMTMR dye (Invitrogen, Carlsbad, CA) for 10 min at 37 °C and adoptively transferred into NOD.SCID.β2 or NOD.SCID mice by tail-vein or retro-orbital injection.

### Two-photon imaging and analysis

Multi-dimensional (x, y, z, time, emission wavelength) two-photon microscopy was employed to image fluorescently labeled lymphocytes in explanted mouse lymph nodes, using a 780 nm femtosecond pulsed laser as described previously (Matheu, Su et al. 2012). Fluorescence emission was split by 510 nm & 560 nm dichroic mirrors into three detector channels, used to visualize eGFP-Orai1E106A transfected (green) adoptively-transferred cells, CMTMR-labelled control cells (red), and collagen (blue). For imaging, lymph nodes were oriented with the hilum away from the water dipping microscope objective (Olympus 20x, NA 0.9) on an upright microscope (Olympus BX51). The node was maintained at 36-37 °C by perfusion with medium (RPMI) bubbled with carbogen (95% O_2_ / 5% CO_2_). 3D image stacks of x=200 μm, y=162 μm, and z=50 μm were sequentially acquired at 18-20 second intervals using MetaMorph software (Molecular Devices, Sunnyvale, CA). This volume collection was repeated for up to 40 min to create a 4D data set. Data were processed and analyzed using Imaris software (Bitplane USA, Concord, MA), and the x,y,z coordinates of individual lymphocytes in the intact lymphoid organ were used to create individual cell tracks.

### Microchannel fabrication and imaging

Microchannel fluidic devices were fabricated by a soft lithography technique with PDMS (polydimethylsiloxane; Sylgard Elastomer 184 kit; Dow Corning, Auburn, MI) as described (Jacobelli, Friedman et al. 2010, Gerard, Patino-Lopez et al. 2014). PDMS base and curing agent were mixed 10:1 and poured onto the silicon master, then left overnight in vacuum. Once the PDMS was set, it was baked at 55 °C for 1 hr and cooled at room temperature. The embedded microchambers were then cut from the mold, and a cell well was punched adjacent to entry into the channels. The PDMS cast and a chambered coverglass (Nunc Lab-Tek, ThermoFisher, Grand Island, NY) were activated for two minutes in a plasma cleaner (Harrick Plasma, Ithaca, NY), bonded together, then incubated at 55 °C for 10 min. Prepared chambers were stored for up to 1 month before use. Prior to imaging, microchambers placed in the plasma cleaner for 5 min under vacuum and 1 min of activation, then coated with 5 µg/mL recombinant human ICAM-1/CD54 Fc (R&D Systems, Minneapolis, MN) in PBS for at least 1 hr at 37 °C. The microchambers were then washed three times with PBS, and T cells were loaded into cell wells (3-5×10^5^ cells resuspended in 10 μL) and incubated at 37 °C for at least 1 hr before imaging.

### Confocal imaging and analysis

Two different Olympus confocal microscopy systems were used to image T cells *in vitro*. For experiments tracking T cell motility in microchambers, we used the self-contained Olympus Fluoview FV10i-LIV, with a 473 nm diode laser for excitation and a 60x phase contrast water immersion objective (NA 1.2). The FV10i-LIV contains a built-in incubator set to 37 °C, together with a Tokai-Hit stagetop incubator to maintain local temperature and humidity. T cells were imaged in RMPI adjusted to 2 mM Ca^2+^ and 2% FCS, and mounted at least half an hour before imaging to allow for equilibration. Cells were imaged at 20-sec intervals for 20-30 min, and the data analyzed using Imaris software. For Ca^2+^ imaging of Salsa6f transfected T cells, we used a Fluoview FV3000RS confocal laser scanning microscope, equipped with high speed resonance scanner and the IX3-ZDC2 Z-drift compensator. Diode lasers (488 and 561 nm) were used for excitation, and two high sensitivity cooled GaAsP PMTs were used for detection. Cells were imaged using the Olympus 40x silicone oil objective (NA 1.25), by taking 4 slice z-stacks at 1.5 μm/step, at 3 sec intervals, for up to 20 min. Temperature, humidity, and CO_2_ were maintained using a Tokai-Hit WSKM-F1 stagetop incubator. Data were processed and analyzed using Imaris software.

### GECI Screening and Salsa6f Plasmid Generation

Plasmids encoding GECIs (GECO and GCaMP6) were obtained from Addgene for screening in live cells. Each probe was cotransfected with Orai1 and STIM1 into HEK 293A cells using Lipofectamine 2000 (Invitrogen, Carlsbad, CA) for 48 hr before screening on an epifluorescence microscope. For construction of Salsa6f, tdTomato was obtained from Addgene, and the pEGP-N1 vector (Clontech, Mountain View, CA) was used as a backbone. GCaMP6f was amplified via PCR with N- and C-terminal primers (5’ CACAACCGGTCGCCACCATGGTCGACTCATCACGTC 3’ and 5’ AGTCGCGGCCGCTTTAAAGCTTCGCTGTCATCATTTGTAC 3’) and ligated into pEGFP-N1 at the AgeI/NotI sites to replace the eGFP gene, while tdTomato was amplified via PCR with N- and C-terminal primers (5’ ATCCGCTAGCGCTACCGGTCGCC 3’ and 5’ TAACGAGATCTGCTTGTACAGCTCGTCCATGCC 3’) and ligated into the backbone at the NheI/BglII sites. An oligo containing the V5 epitope tag was synthesized with sense and antisense strands (5’ GATCTCGGGTAAGCCTATCCCTAACCCTCTCCTCGGTCTCGATTCTACG 3’ and 5’ GATCCGTAGAATCGAGACCGAGGAGAGGGTTAGGGATAGGCTTACCCGA 3’) and ligated into the backbone at the BglII/BamHI sites, linking tdTomato to GCaMP6f and creating Salsa6f. The amplified regions of the construct were verified by sequencing (Eton Bioscience Inc., San Diego, CA). This plasmid, driven by the CMV promoter, was used for transient transfections in HEK 293A cells with Lipofectamine 2000 and in primary human T cells with Amaxa Nucleofection.

## Data Analysis and Statistical Testing

Samples sizes were comparable to previous single cell analyses of motility (Jacobelli, Friedman et al. 2010, Greenberg, Yu et al. 2013, Gerard, Patino-Lopez et al. 2014). Each experiment used separate isolations of human T cells from different donors. With the exception of momentary velocities in Figure 6, each measurement corresponds to a different cell. Mean ± standard error of the mean was used as a measure of the central tendency of distributions. Video analysis was performed using Imaris software, Spots analysis was used for tracking of cell velocity and Volumes analysis was used for measuring total fluorescence intensity of GECI probes. To reduce selection bias in our analysis of motility and trajectory, all clearly visible and live cells were tracked from each video segment. The arrest coefficient is defined as fraction of time each cell had an instantaneous velocity < 2 μm/min. The coefficient of variation was defined for each individual cell as the standard deviation divided by the mean of its instantaneous velocity. For Salsa6f imaging analysis, ratio (R) was calculated by total GCaMP6f intensity divided by total tdTomato intensity, while initial ratio (R_0_) was calculated by averaging the ratios of the first five time points in each individual cell trace. Photobleaching of tdTomato fluorescence intensity (20-30% decline) was corrected in ratio calculations, as a linear function of time. Figures were generated using Prism 6 (GraphPad Software, San Diego, CA) and Origin 5 (OriginLabs, Northampton, MA). Due to the expectation that individual cells exhibit multiple motility modes, and to avoid assumptions concerning the shapes of motility distributions, non-parametric statistical testing was performed (Mann-Whitney *U* test, unpaired samples, two-tailed). Differences with a *P* value of ≤ 0.05 were considered significant: \**P* ≤ 0.05; \*\**P* < 0.01; \*\*\**P* < 0.005; \*\*\*\**P* < 0.001. Similar distributions were compared using the Hodges-Lehmann median difference value and 95% confidence intervals under the assumption that the starting distributions had similar shapes.

## Supplementary Information

**Supplementary Figure 1.**
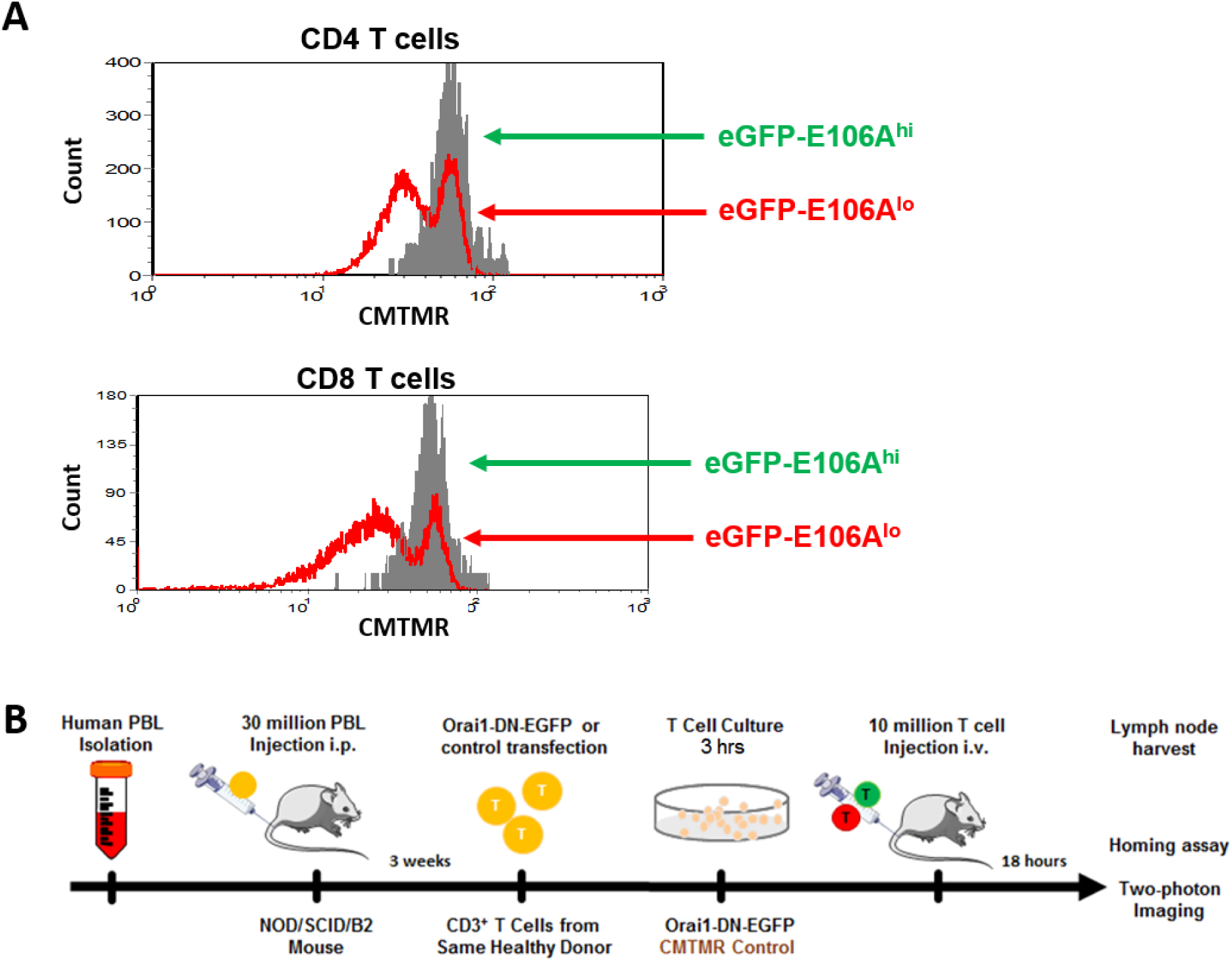
Validation of dominant-negative eGFP-Orai1-E106A to assess activation, homing, and motility of human T cells. (**A**) Primary human CD4^+^ and CD8^+^ T cells were transfected with eGFP-E106A, then uniformly labeled with the fluorescent cell tracker dye CMTMR and co-cultured with SEB-pulsed primary human dendritic cells from the same donor; proliferation was assessed after 72 hr by CMTMR dilution as measured by flow cytometry. (**B**) Protocol for homing and two-photon imaging of transfected human CD3^+^ T cells in reconstituted NOD.SCID.β2 mouse lymph node.

**Video 1**. HEK 293A cells transfected with Salsa6f, first washed with 0 mM Ca^2+^ followed by 2 μΜ ionomycin in 2 mM Ca^2+^; scale bar = 20 μm, time shown in hr:min:sec.

**Video 2**. Salsa6f transfected human T cell in confined microchannel, with merged red (tdTomato), green (GCaMP6f), and DIC channels; scale bar = 10 μm, time shown in hr:min:sec.

**Video 3**. Salsa6f transfected human T cells in open microchamber, with merged red (tdTomato), green (GCaMP6f), and DIC channels, circular structures are support pillars part of the PDMS microchamber; scale bar = 10 μm, time shown in hr:min:sec.

